# Beyond signaling activation: Phosphorylation modulates Grb2 phase separation to create multivalent scaffolds

**DOI:** 10.64898/2026.03.07.710134

**Authors:** Raphael Vinicius R. Dias, Carolina G. Oliveira, Aline Sebastiane de Oliveira, Guilherme Dias Fusari, Sebastião Roberto Taboga, Antonio José Costa-Filho, Fernando A. de Melo

## Abstract

The Growth Factor Receptor-Bound Protein 2 (Grb2) is a central adaptor protein in signal transduction pathways, yet how its monomer-dimer equilibrium governs its supramolecular organization remains elusive. Here, we demonstrate that the oligomeric state of Grb2 acts as a binary switch for Liquid-Liquid Phase Separation (LLPS). While the auto-inhibited homodimer forms only transient, thermodynamically unstable assemblies in crowding conditions, the phosphorylation-mimetic monomer (Y160E) drives the formation of robust, gel-like condensates. Integrating turbidity assays, temperature-controlled dynamic light scattering, fluorescence recovery after photobleaching (FRAP), hyperspectral imaging analyses, and coarse-grained molecular dynamics simulations, we reveal that this phase transition is enthalpy-driven and reliant on a specific electrostatic network between the SH2 domain residue R142 and a C-terminal SH3 acidic cluster (Q170-D172). Crucially, we show that these monomeric condensates function as "scaffolds" that efficiently recruit and sequester cytosolic wild-type dimers ("clients") into the dense phase. This recruitment mechanism resolves the paradox of how non-condensing wild-type proteins participate in phase separation. Our findings propose a novel regulatory model where phosphorylation nucleates the formation of high-density signaling hubs, redefining Grb2 from a passive adaptor to a dynamic spatial organizer of the Ras/MAPK pathway.

## Introduction

The Growth Factor Receptor-bound Protein 2 (Grb2) is a ubiquitous and critical adaptor protein that bridges activated cell-surface receptors, such as Receptor Tyrosine Kinases (RTKs), to downstream intracellular signaling pathways, most notably the RAS/MAPK cascade [1, 2]. Structurally, Grb2 is composed of a central Src Homology 2 (SH2) domain flanked by two Src Homology 3 (SH3) domains (SH3-SH2-SH3), connected by flexible linkers that allow significant inter-domain plasticity **(Figure 1)** [3]. This modular architecture enables Grb2 to function as a cross-linker: the SH2 domain binds to specific phosphotyrosine motifs on receptors (e.g., EGFR), while the SH3 domains constitutively bind proline-rich motifs on effectors such as SOS1 and Gab2 [4, 5]. Through these interactions, Grb2 regulates vital cellular processes including proliferation, differentiation, and survival, and its dysregulation is frequently implicated in oncogenesis [6, 7].

**Figure 1.**
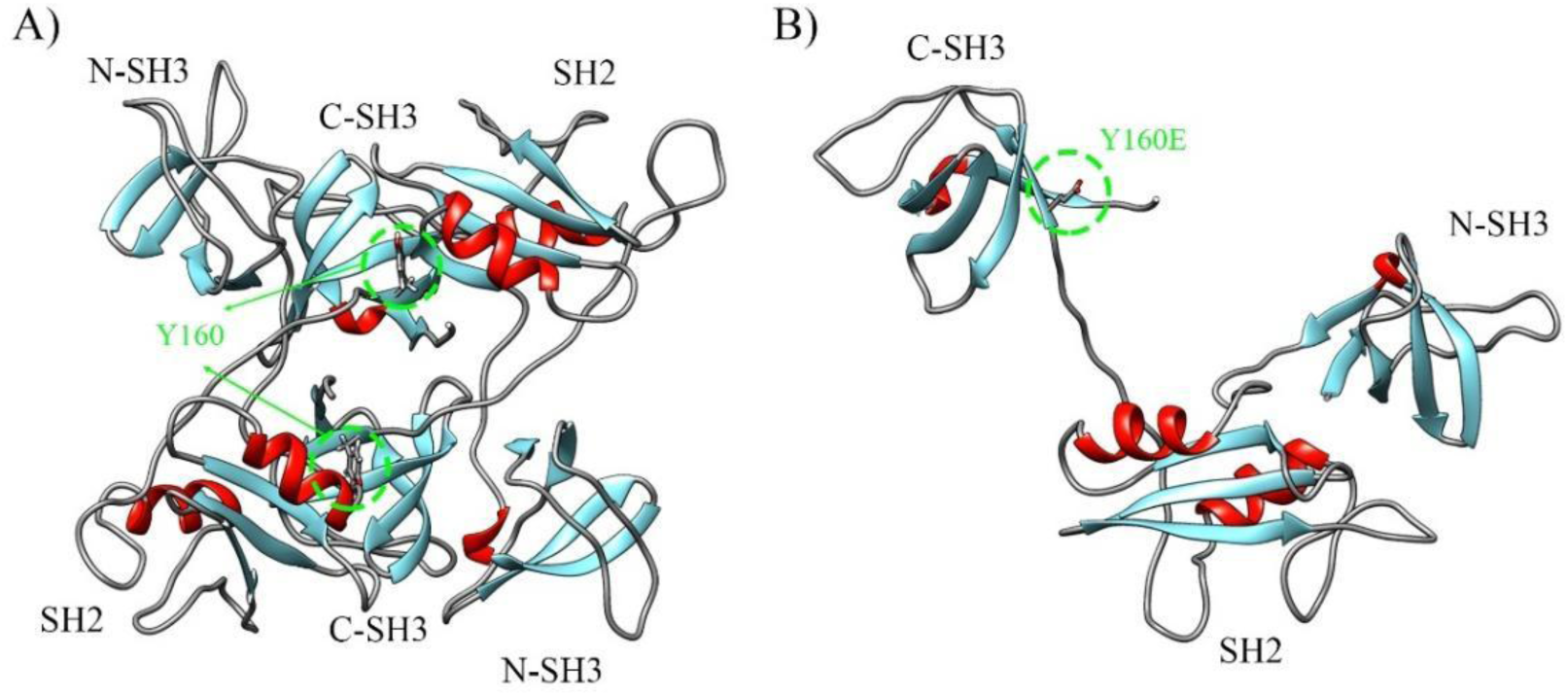
Structural reorganization of Grb2 triggered by the phosphomimetic mutation Y160E. **(A)** Ribbon representation of the Wild-Type Grb2 homodimer (PDB:1GRI). The protein adopts a closed, auto-inhibited configuration stabilized by intermolecular contact between the SH2 and C-terminal SH3 domains. The key Tyrosine 160 (Y160) residue, located at the dimerization interface, is highlighted in green. **(B)** Structural model of the Grb2 Y160E mutant predicted by AlphaFold 3. The introduction of a negative charge (glutamate) at position 160 mimics phosphorylation, disrupting the dimer interface and releasing the protein into an open, highly flexible monomeric state. Structures are colored by secondary structure elements (α-helices in red, β-sheets in cyan, and loops/coils in grey), highlighting the modular architecture and the exposure of binding domains in the monomeric form.

While often depicted as a simple adaptor, the biophysical behavior of Grb2 is strictly regulated by a dynamic equilibrium between monomeric and dimeric states. In the cytoplasm, Grb2 predominantly exists as a homodimer, stabilized by intermolecular interactions involving the SH2 and C-terminal SH3 domains [8]. This dimeric configuration functions as an auto-inhibited state that imposes a threshold on signaling. Lin et al. demonstrated that dimeric Grb2 binds to the C-terminus of FGFR2, forming a latent complex that prevents inadvertent pathway activation [9]. The transition to a functional state requires the dissociation of this dimer, triggered by phosphorylation at Tyrosine 160 (Y160) or by high-affinity ligand binding,which releases Grb2 to engage downstream effectors such as SOS1, thereby driving the RAS/MAPK cascade [8].

Recent advances in cell biology have revealed that signal transduction is not merely governed by diffusion-limited collisions but is spatially organized within membraneless organelles formed via Liquid-Liquid Phase Separation (LLPS) [10, 11]. These biomolecular condensates concentrate enzymes and substrates, enhancing reaction kinetics and specificity while sequestering components from the surrounding cytosol [10]. The formation of these condensates is typically driven by multivalent interactions involving modular domains (such as SH2 and SH3) and Intrinsically Disordered Regions (IDRs). Li et al. demonstrated that multivalent SH3-PRM systems undergo sharp phase transitions [12]. Consistent with this, Grb2 has been identified as a key driver of phase separation when cross-linked by multivalent partners, such as SOS1 or phosphorylated scaffolds, in T-cell receptor signaling [13]. However, while recent structural studies have hinted at the potential of Grb2 to self-assemble into meshworks **[15]**, the thermodynamic determinants governing this process remain poorly understood. Furthermore, resolving the material nature of these assemblies, whether they behave as dynamic liquids or arrested gels, is a critical open question, as the physical state of a condensate directly dictates its function and reversibility in a cellular context **[14]**.

In this study, we expand upon the regulatory framework established by Ahmed et al., who demonstrated that the monomer-dimer equilibrium acts as a critical checkpoint for signaling fidelity, with the dimer functioning as an auto-inhibited state [8]. Here, we define this oligomeric switch as a master regulator of Grb2 phase behavior. Focusing on the phosphorylation-mimetic mutant Y160E (constitutively monomeric) versus the Wild-Type protein, we combine turbidity assays, microscopy images, and Coarse-Grained Molecular Dynamics (CG-MD) simulations to dissect the molecular mechanism of condensation. We demonstrate that the "uncaging" of the monomer exposes a specific electrostatic network, providing a thermodynamic basis for the self-assembly potential suggested by Breithaupt et al. **[15]**. Furthermore, we characterize the resulting condensates as stable, gel-like materials **[14]**, capable of recruiting cytosolic WT dimers. Our findings reveal a "Scaffold-Client" mechanism [16] that resolves how the non-condensing WT protein participates in phase separation, proposing a new model for spatial organization in the RAS/MAPK pathway.

## Results and Discussion

### The monomeric state of Grb2 (Y160E) drives robust condensate formation, whereas the dimeric WT forms transient, thermodynamically unstable assemblies

To dissect the contribution of the oligomeric state to Grb2 supramolecular organization, we first mapped the phase behavior of the constitutive monomer Y160E. Using high-throughput turbidity assays across a range of protein (20–100 µM) and crowding agent concentrations (0–20% PEG^6000^), we systematically established the boundaries for phase separation. As shown in **Figure 2**, Grb2 Y160E exhibited a distinct phase envelope, with a critical saturation concentration (C_sat_) established around 80 µM. At 100 µM, turbidity peaked between 14% and 16% PEG, followed by a slight decrease at 20% PEG. This re-entrant behavior suggests an optimal crowding window where attractive intermolecular forces are maximized relative to excluded-volume effects. Representative microscopy (**Figure 2, inset**) confirmed that these turbidity signals correspond to the formation of abundant, spherical liquid droplets, characteristic of condensate formation .

**Figure 2.**
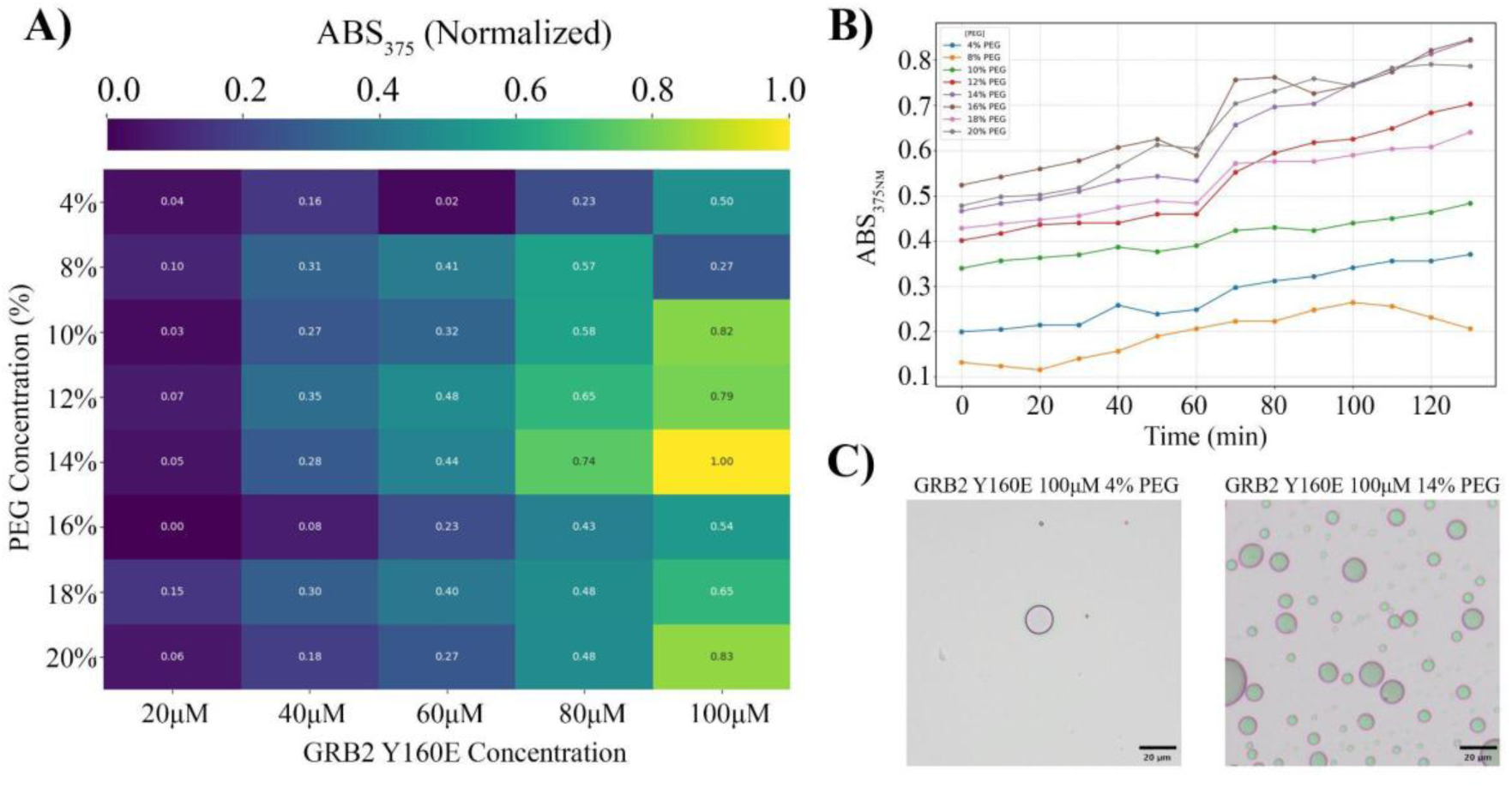
Phase behavior and liquid-like nature of the monomeric Grb2 Y160E. Phase diagram derived from solution turbidity measurements (ABS_375_) as a function of protein concentration (20–100 µM) and PEG^6000^ (4–20%). The heatmap represents normalized turbidity values ranging from 0 (dark purple, dispersed phase) to 1.0 (bright yellow, maximum condensation). Data were normalized relative to the peak absorbance observed at the condition of 100 µM protein and 14% PEG^6000^. Representative Differential Interference Contrast (DIC) microscopy images (inset) confirm the formation of spherical liquid droplets at 100 µM protein in the presence of 4 and 14% PEG^6000^.

Thermodynamic stability is not intrinsic to all Grb2 species. When we compared the temporal evolution of those droplets with that of the dimeric Wild-Type (WT) protein under maximal crowding conditions (Supplementary Fig. S1), we revealed a striking kinetic dichotomy (**Figure 3**). The WT protein (red trace) underwent a rapid initial condensation driven by crowding forces but failed to sustain the phase-separated state, exhibiting a sharp exponential decay with droplets dissolving almost completely within 60 minutes. In contrast, the monomeric Y160E (black trace) displayed a robust, stepwise growth profile that stabilized at high turbidity values.

**Figure 3.**
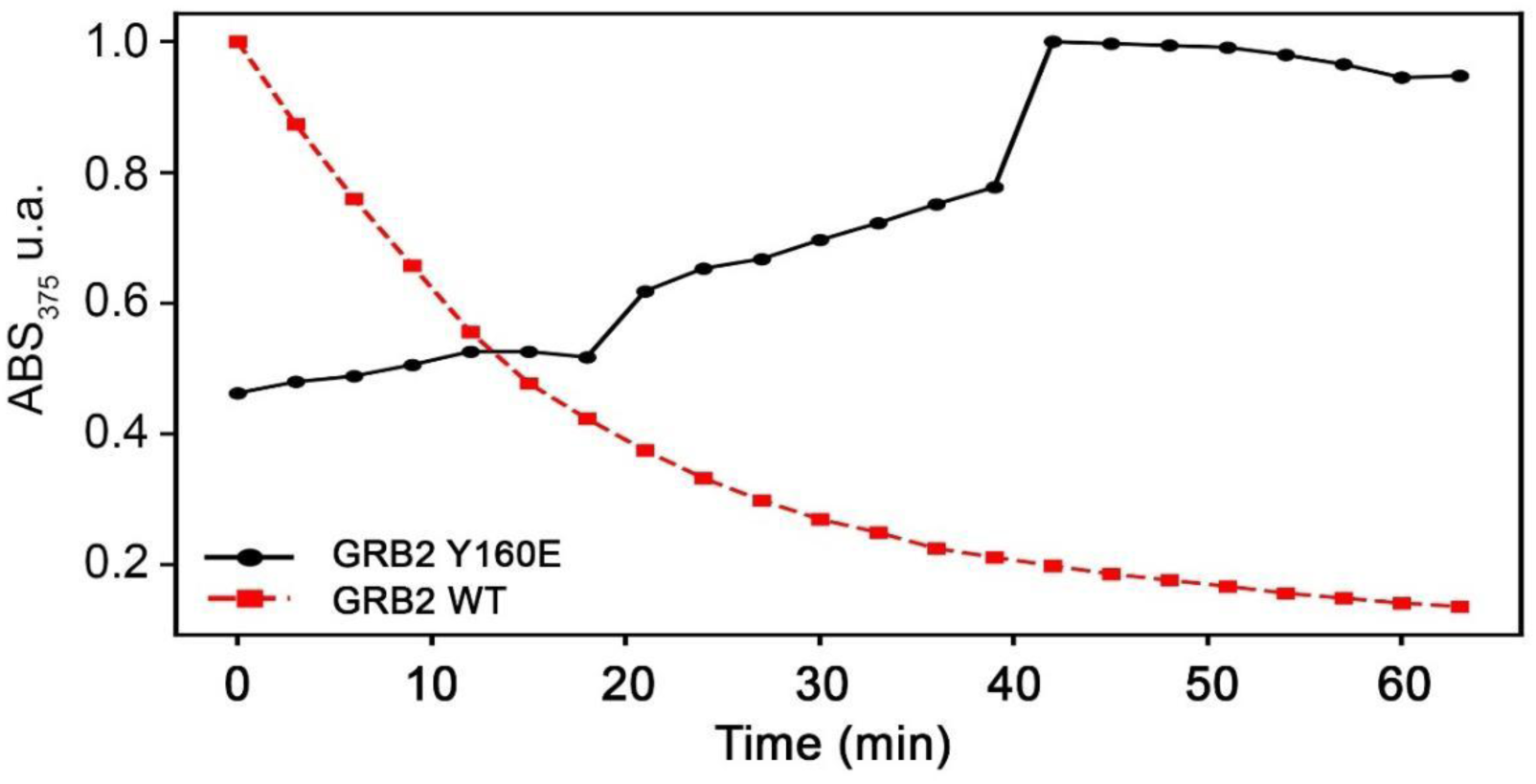
Oligomerization state dictates the kinetic stability of Grb2 condensates. Comparative kinetic analysis of condensate formation monitored by turbidity (ABS_375_) over time. The assays were performed with Grb2 Y160E (black circles) and Wild-Type (red squares) under identical crowding conditions (100 µM protein and 20% PEG^6000^). While the monomeric Y160E exhibits a robust and progressive increase in turbidity consistent with stable condensate maturation, the dimeric WT displays a rapid exponential decay (t_1/2_ ∼ 10 min), indicating the transient and thermodynamically unstable nature of its phase separation.

These data suggest that the dimeric configuration functions as a **"**kinetic trap" that disfavors phase separation. We propose that while crowding agents can transiently overcome the entropic barrier to induce nucleation of WT dimers, the autoinhibited SH2/SH3 interface prevents the formation of the stable percolated network required for droplet maturation. Conversely, the Y160E mutation "uncages" these domains, allowing the monomer to act as a competent scaffold for persistent, thermodynamically stable condensation.

### Phase separation is driven by specific electrostatic stickers and exhibits thermal reversibility

To rigorously validate the phase boundaries observed in our initial screening and confirm the specific nature of the condensates, we performed a refined titration using the crowding agent PEG^6000^ (0–22%). As shown in Figure 4A, turbidity measurements revealed a sharp sigmoidal phase transition, with the onset of condensation at 8% PEG and a clear saturation plateau between 16% and 18%. To confirm that this turbidity arises from genuine protein sequestration rather than non-specific aggregation, we used fluorescence microscopy with RED-NHS-labeled Grb2 Y160E (Figure 4B). The imaging revealed distinct, micron-sized spherical droplets with high fluorescence intensity, confirming that the monomeric protein efficiently partitions into the condensed phase, a hallmark of Liquid-Liquid Phase Separation (LLPS).

**Figure 4.**
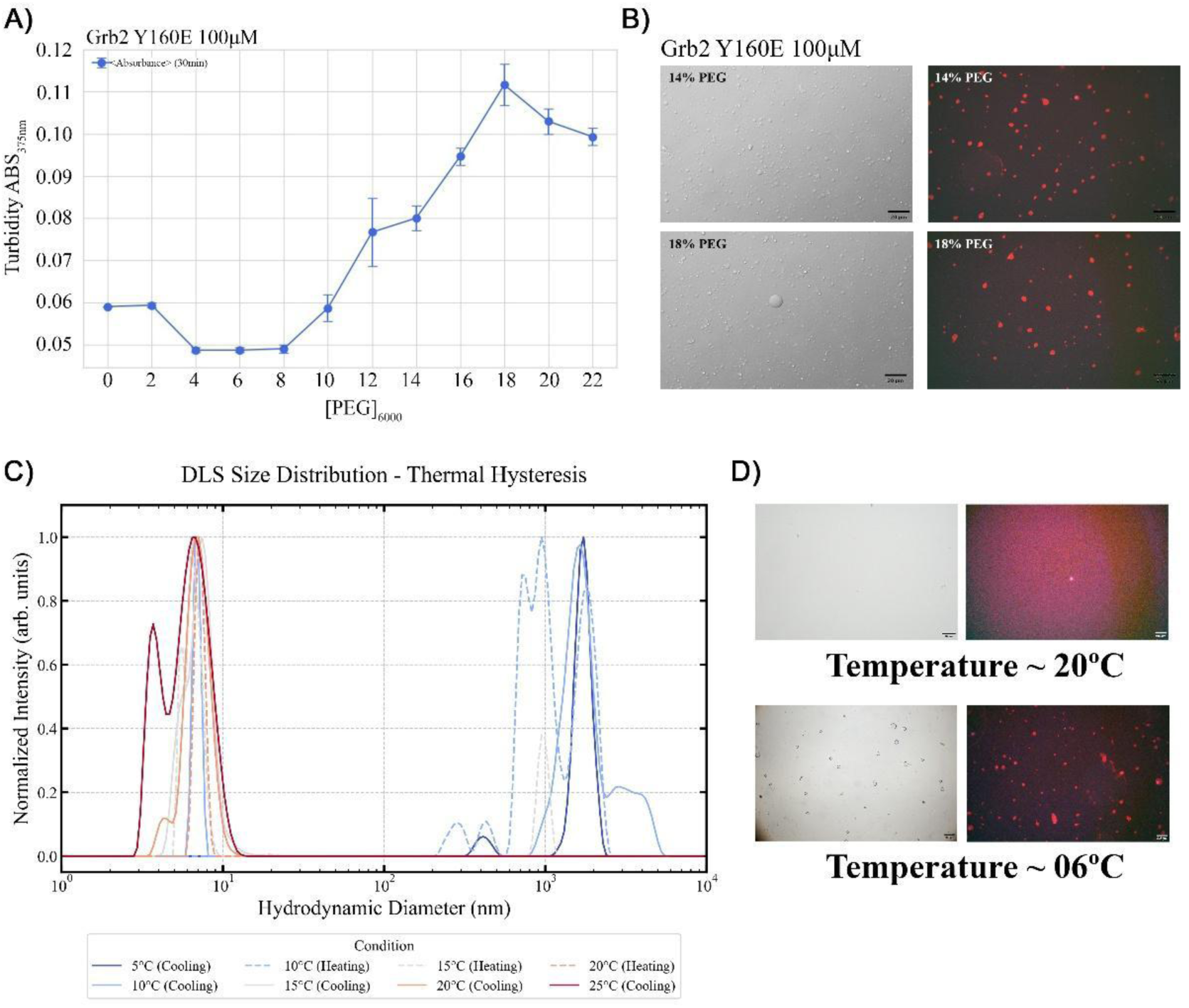
Grb2 Y160E undergoes reversible liquid-liquid phase separation driven by crowding and temperature. **(A)** Detailed phase diagram derived from turbidity (ABS_375_) titration of PEG^6000^ (0–22%) with Grb2 Y160E fixed at 100 µM. Error bars represent the standard deviation of triplicates. **(B)** Differential Interference Contrast (DIC) and Fluorescence microscopy of RED-NHS labeled Grb2 Y160E at 14% and 18% PEG, confirming the sequestration of Grb2 into liquid droplets. **(C)** Dynamic Light Scattering (DLS) analysis of the thermal response. Size distribution profiles show a reversible transition from a monodisperse population (∼5 nm) at 20°C (red/orange curves) to large mesoscopic assemblies (>1000 nm) at 5°C (blue curves), exhibiting thermal hysteresis. **(D)** Microscopy validation of thermally induced phase separation. The protein solution appears homogeneous at 20°C but spontaneously forms spherical condensates upon cooling to 6°C in the absence of crowding agents.

To determine whether this phase separation was solely dependent on crowding agents or represented an intrinsic property of the monomeric protein, we investigated its behavior using temperature as a control variable in the absence of PEG. We monitored the hydrodynamic diameter (D_h_) using Dynamic Light Scattering (DLS) across a cooling-heating cycle (20°C – 5°C – 20°C).

At room temperature (20°C), the protein existed predominantly as a monodisperse population with a hydrodynamic diameter of ∼5 nm (red curves, Figure 4C). However, cooling the sample below a critical temperature (Tc ∼10°C) triggered a dramatic phase transition, shifting the distribution towards large mesoscopic assemblies (> 1000 nm) (blue curves). Strikingly, this transition exhibited thermal hysteresis but was fully reversible; upon re-heating to 20°C, the condensates dissolved, and the monomeric population recovered. This reversibility, visually corroborated by the spontaneous formation of droplets at ∼6°C (Figure 4D), distinguishes these assemblies from the irreversible, pathological fibrillization often observed in neurodegenerative disorders **[14]**, identifying the condensates as dynamic, metastable states.

The observation that condensation is triggered by cooling (Upper Critical Solution Temperature, UCST-like behavior) provides critical insights into the thermodynamics of the system. It implies that the process is enthalpy-driven, where the favorable enthalpy change (ΔH < 0) arising from specific intermolecular interactions overcomes the entropic penalty (TΔS) of organizing monomers into droplets. The observed hysteresis loop suggests the existence of a nucleation barrier, consistent with a first-order phase transition [22]. We propose that this behavior is potentiated by the Y160E mutation itself. By introducing a negative charge to mimic phosphorylation, the mutation not only disrupts the autoinhibited dimer interface to release the monomer but likely increases the conformational flexibility of the protein. This "uncaging" effect facilitates the exposure of critical contact sites, such as arginine/lysine patches necessary for the electrostatic networks that drive this enthalpy-dependent condensation.

### Coarse-grained simulations reveal an electrostatic lock-and-key mechanism between SH2 and SH3 domains

To decipher the molecular origins of the enthalpy-driven condensation observed experimentally, we performed Molecular Dynamics (MD) simulations using the residue-level Coarse-Grained (CG) force field CoCoMo [18]. This approach allowed us to simulate the spatiotemporal evolution of large protein ensembles at concentrations comparable to our experimental setup, bridging the gap between discrete molecular interactions and mesoscopic phase behavior.

The simulation trajectory successfully captured the spontaneous phase separation of Grb2 Y160E (**Figure 5A**). Starting from a dispersed monomeric state, the system evolved through a nucleation phase, forming small transient clusters which subsequently coalesced into a single, dense, and stable condensate (t = 500 ns). Notably, the kinetics of cluster growth observed in silico closely mirrored the saturation profile measured in our turbidity assays (**Figure 3**), validating the ability of the computational model to capture the essential physics of the phase transition.

**Figure 5.**
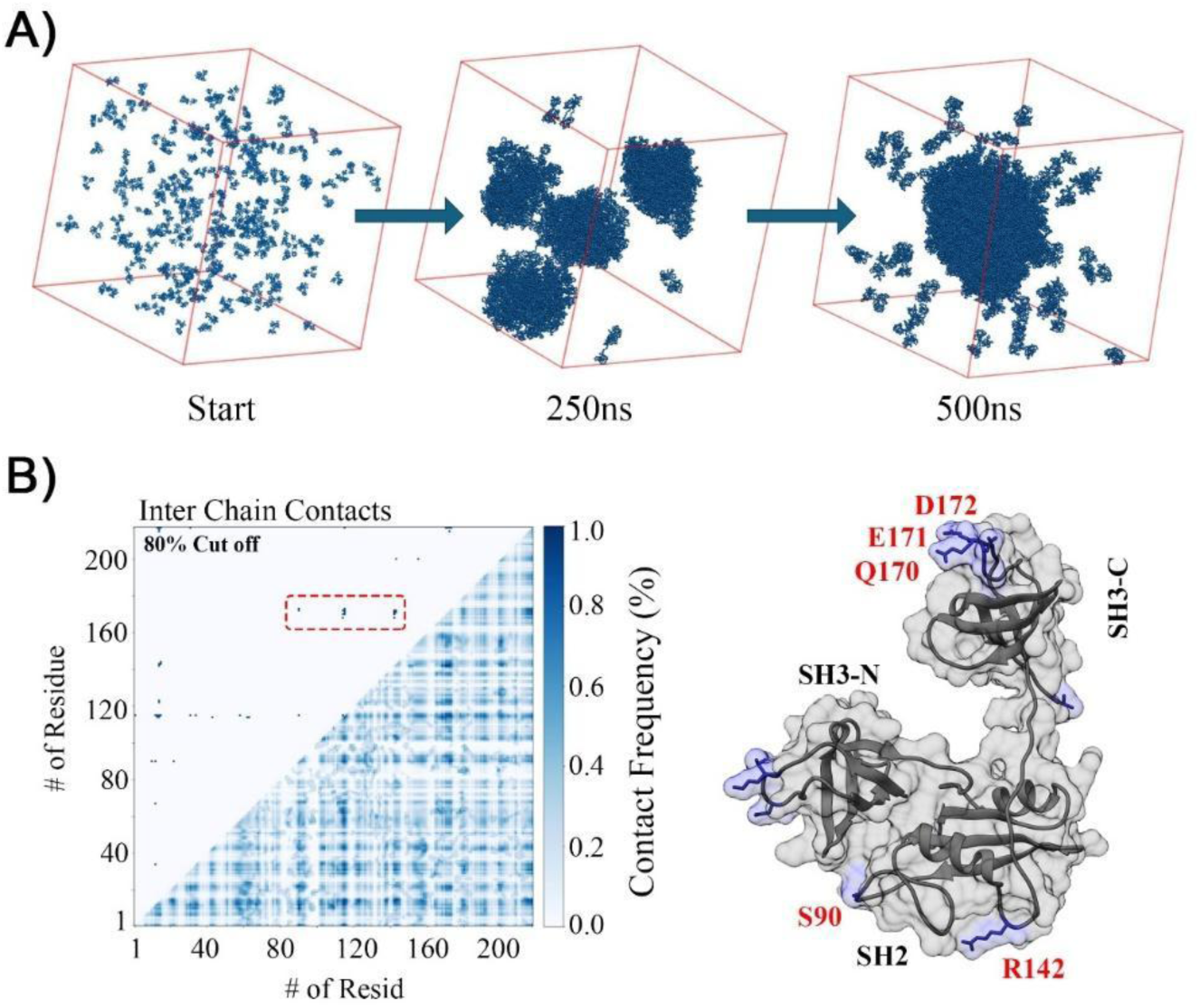
Molecular mechanism of Grb2 Y160E phase separation. **(A)** Snapshots from Coarse-Grained Molecular Dynamics (CG-MD) simulations showing the spontaneous condensation of Grb2 Y160E. The system evolves from a dispersed solution (Start) to formed clusters (250 ns) and finally a stable, single condensate (500 ns). **(B)** *Left:* Intermolecular contact map calculated from the condensed phase trajectory (80% cutoff). The red box highlights the dominant off-diagonal interaction between the SH2 domain (residues 60-150) and the C-terminal SH3 domain (residues 160-217). *Right:* Structural representation of the key "sticker" residues identified in the contact map. The positively charged R142 (SH2 domain) and the negatively charged cluster Q170/E171/D172 (SH3-C domain) are highlighted as the primary electrostatic drivers of the intermolecular network.

We next analyzed the intermolecular contact frequencies within the condensed phase to identify the stickers driving this assembly. The contact map (Figure 5B, left) revealed a non-random interaction pattern, strictly dominated by a specific interface between the central SH2 domain and the C-terminal SH3 domain (SH3-C). Mapping these high-frequency contacts onto the Grb2 structure highlighted a critical electrostatic interface (Figure 5B, right): the positively charged Arginine 142 (R142) in the SH2 domain engages in stable salt-bridge-like interactions with a negatively charged acidic cluster comprising Q170, E171, and D172 in the SH3-C domain.

This molecular mechanism provides a structural rationale for the self-assembly properties recently reported by **Breithaupt et al. [15]**. Their study demonstrated that Grb2 constructs lacking the C-terminal SH3 domain fail to form meshworks, whereas the SH2-SH3C module is sufficient for assembly. Our simulations clarify that this loss of function arises specifically from the deletion of the acidic sticker cluster (Q170-D172). By removing these residues, the truncation breaks the electrostatic binary code required for networking. Thus, the open monomeric conformation of Y160E is essential not only to uncage the SH2 domain but to expose its complementary SH3-C partner, generating the favorable enthalpy required to drive condensation.

### Grb2 Y160E condensates exhibit gel-like material properties and scaffold the recruitment of wild-type dimers

Our Coarse-Grained simulations predicted that the "uncaged" monomeric interface drives the formation of a robust, high-connectivity electrostatic network. To experimentally probe the material state that emerges from these interactions, we combined fluorescence recovery after photobleaching (FRAP) with hyperspectral imaging (HSI) and phasor-plot analysis to map polarity and molecular crowding within Grb2 condensates. For HSI, we used ACDAN (6-acetyl-2-(dimethylamino)naphthalene), a solvatochromic probe widely employed to report dipolar relaxation in cells, tissues, and biomimetic systems [28]. Upon excitation, ACDAN undergoes a change in dipole moment: in water-rich, highly fluid environments, efficient solvent relaxation produces a red-shifted emission, whereas in crowded, less fluid microenvironments, where relaxation is hindered, its emission shifts to the blue [29]. Using this integrated approach, we first assessed the exchange dynamics of Grb2 Y160E condensates by FRAP (Figure 6A). Consistent with the formation of a percolated network, the condensates exhibited minimal fluorescence recovery (< 10%) over the monitored time course (>120s). This slow exchange dynamic indicates that the molecules within the droplet are not freely diffusing as in a simple Newtonian liquid, but are kinetically arrested in a gel-like or glassy state. While liquid-to-solid transitions are often associated with pathological aggregation in neurodegenerative proteins [14], here we propose that this physical hardening serves a functional purpose: it stabilizes the condensate against dissolution, creating a persistent structural platform. To further distinguish this kinetically arrested state from non-specific amorphous aggregation, we employed hyperspectral imaging coupled with spectral phasor (phasor-plot) analysis (Supplementary Figure S2). This approach revealed a single, compact spectral population and spatially uniform phasor maps within Grb2 Y160E droplets, consistent with a homogeneous microenvironment and a well-defined dense phase rather than heterogeneous amorphous aggregates.

**Figure 6.**
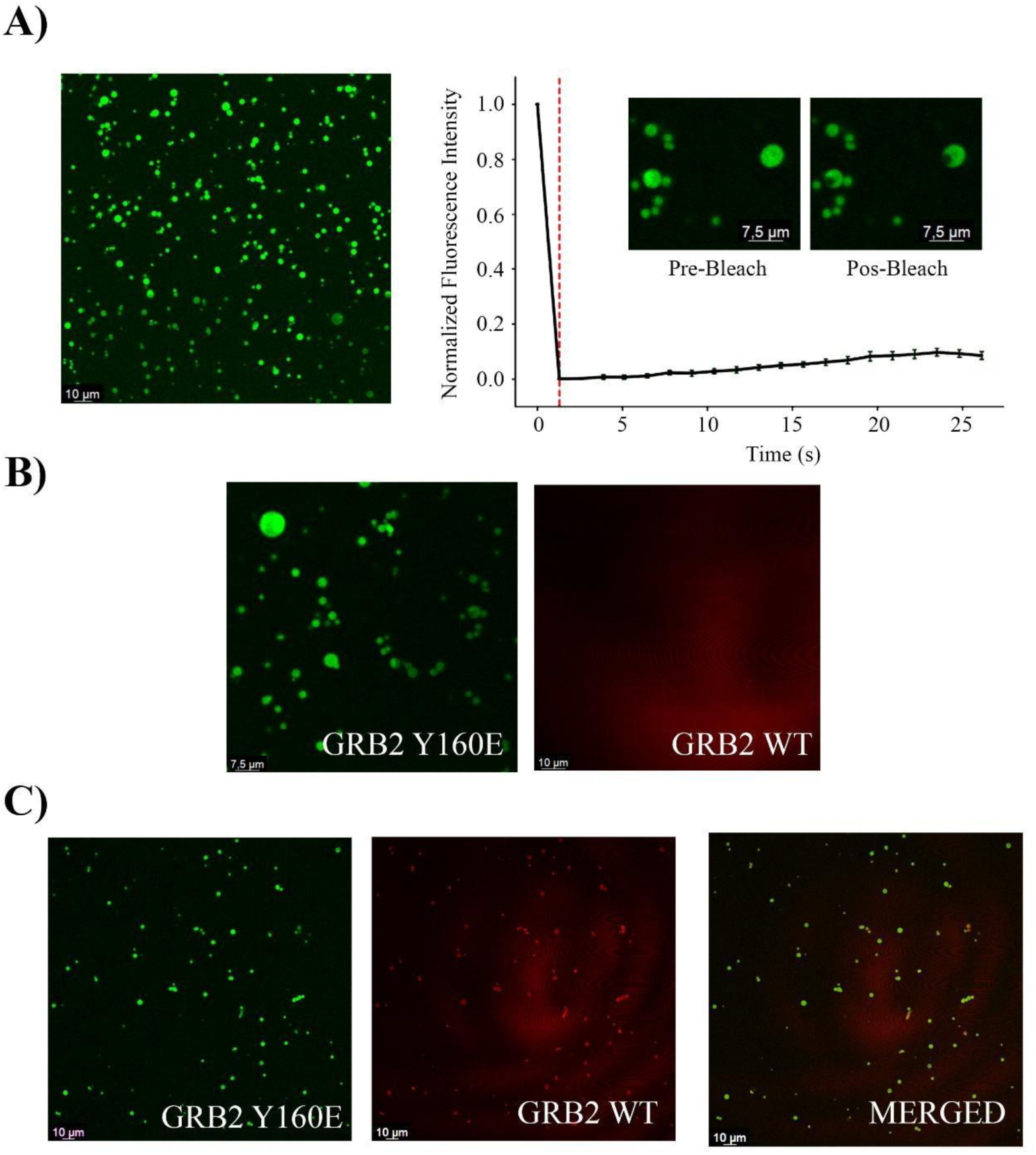
Material properties and co-condensation of Grb2 variants. (A) Fluorescence Recovery After Photobleaching (FRAP) analysis of Grb2 Y160E condensates (Green). Left: Representative confocal image pre-bleach. Right: Normalized fluorescence recovery curve. The lack of significant recovery indicates a gel-like material state with restricted molecular diffusion. Insets show the bleached region at different time points. (B) Confocal microscopy controls showing that Grb2 Y160E (Green) forms droplets autonomously, whereas Grb2 WT (Red) remains dispersed under identical conditions (100 µM protein, 14% PEG-6000). (C) Co-condensation assay. Mixing Grb2 Y160E (Green) and Grb2 WT (Red) results in the formation of condensates containing both proteins. The merged image (Yellow) demonstrates the efficient recruitment (partitioning) of the WT "client" into the "scaffold" formed by Y160E.

We next investigated whether this stable scaffold formed by the activated monomer could influence the behavior of the latent dimeric WT protein. Using the Scaffold-Client framework, in which a phase-separating protein (scaffold) recruits non-phase-separating partners (clients) **[10],** we designed a co-condensation assay. As established previously, Grb2 WT labeled with a red fluorophore failed to form condensates on its own under identical conditions, appearing as a diffuse background (**Figure 6B**).

Then, when we mixed the scaffold-forming Y160E (Green) with the client WT (Red), we observed a striking recruitment phenomenon. As shown in **Figure 6C**, the WT protein was efficiently partitioned into the dense phase generated by Y160E, resulting in droplets with strong colocalization of both fluorophores. This result confirms that while the WT dimer lacks the valency to nucleate phase separation (acts as a client), it retains sufficient surface interactions to bind the pre-formed Y160E network (the scaffold).

Biologically, this recruitment mechanism suggests a novel layer of spatial regulation for the RAS/MAPK pathway. We propose that phosphorylation at Y160 does more than simply activate Grb2; it acts as a nucleation event. By generating a local population of monomers that self-assemble into stable gels, the cell creates high-density signaling hubs. These hubs can then trap cytosolic WT dimers, potentially functioning as amplification chambers, where high local concentrations facilitate effector binding, or as molecular buffers that sequester Grb2 to prevent spurious signaling noise in the cytosol.

Taken together, our data define a biophysical framework in which the oligomeric state of Grb2 acts as a master regulator of its material phase. We established that the canonical homodimer functions as a kinetic trap, structurally sequestering the electrostatic "stickers" (R142 and D172) required for networking. Phosphorylation, mimicked here by the Y160E mutation, breaks this trap, exposing a binary electrostatic code that drives enthalpy-dependent condensation. Importantly, we demonstrated that the resulting condensates are not merely liquid droplets but mature into viscoelastic scaffolds. This gel-like architecture is mechanically sufficient to recruit cytosolic components, enabling the "Scaffold-Client" mechanism observed with the wild-type protein. Thus, the transition from a transient, unstable assembly (WT) to a stable, recruiting compartment (Y160E) represents a shift in the physical nature of Grb2, equipping the cell with a switchable mechanism to organize signaling molecules in space and time.

## Conclusions

In this study, we established that the oligomerization state of Grb2 acts as a binary switch for its supramolecular organization. While the canonical homodimer functions as a **kinetic trap** due to the autoinhibition of its binding interfaces, the phosphorylation-mimetic monomer (Y160E) is uncaged, revealing a multivalent architecture capable of driving robust Liquid-Liquid Phase Separation (LLPS). Our thermodynamic and computational analyses demonstrate that this process is enthalpy-driven, relying on a specific **electrostatic lock-and-key network** between residue R142 in the SH2 domain and the acidic cluster (D172/E171) in the C-terminal SH3 domain. This identifies the monomeric Grb2 not merely as a signaling unit, but as a structural scaffold programmed to self-associate under physiological conditions.

Critically, we demonstrated that these condensates mature into a gel-like state with restricted molecular diffusion. Distinct from the irreversible fibrillization characteristic of pathological aggregates [14], the Grb2 condensates remain thermally reversible. This kinetic arrest represents a stable material state that allows the monomeric scaffold to sequester and recruit cytosolic WT dimers. This "Scaffold-Client" mechanism resolves the apparent paradox of how the non-condensing WT protein can participate in phase separation without external cross-linkers

Biologically, our findings suggest a novel layer of regulation mechanism for the RAS/MAPK pathway. We propose that phosphorylation at Y160 does more than activate Grb2; it nucleates the formation of high-density signaling hubs. These membraneless organelles could function either as amplification chambers—concentrating effectors to exceed activation thresholds—or as molecular buffers that physically sequester Grb2 dimers to dampen noise. By integrating biophysical characterization with molecular simulation, this work redefines Grb2 into a dynamic spatial organizer, opening new avenues to explore how dysregulation of phase separation contributes to oncogenic signaling.

## Materials and Methods

### Protein Expression and Purification

The DNA sequence encoding full-length human Grb2 (residues 1–217) was cloned into a pET-28a vector containing an N-terminal 6xHis-tag. The Y160E mutation was introduced by site-directed mutagenesis and confirmed by DNA sequencing. Both Wild-Type (WT) and Y160E constructs were transformed into *E. coli* BL21 (DE3) cells for protein expression. Protein expression and purification were performed following the protocol described by **Ahmed et al. [8]**, with minor modifications. Briefly, cultures were grown in Luria-Bertani (LB) medium at 37°C under agitation **(180 rpm)** until an optical density (OD_600_) of 0.7 was reached. Protein expression was induced by the addition of 0.3 mM Isopropyl β-D-1-thiogalactopyranoside (IPTG), and cells were incubated at 20°C for 16 hours.

Cells were harvested by centrifugation (4000 rpm for 20 min at 4°C) and resuspended in Lysis Buffer (50 mM Tris-HCl pH 7.5, 500 mM NaCl, 20 mM Imidazole). Lysis was performed by sonication (20 cycles of 30s on/30s off) in ice. The lysate was clarified by centrifugation (17.000 rpm for 45 min at 4°C), and the supernatant was loaded onto a Ni-NTA affinity column (Qiagen or GE Healthcare) pre-equilibrated with Lysis Buffer. Bound proteins were washed to remove non-specific interactions and eluted with Elution Buffer (50 mM Tris-HCl pH 7.5, 500 mM NaCl, 300 mM Imidazole).

A final polishing step was performed using Size Exclusion Chromatography (SEC) on a Superdex 75 10/300 GL column (Cytiva) equilibrated in Assay Buffer (50 mM HEPES pH 7.4, 300 mM NaCl, 0. Protein purity 5 mM TCEP). (>95%) was assessed by SDS-PAGE, and concentrations were determined by absorbance at 280 nm using the theoretical extinction coefficients calculated via the ProtParam tool on the ExPASy server [17]. The oligomeric state (monomer vs. dimer) was verified by the elution volume in analytical SEC compared to standard molecular weight markers.

### Fluorescence Labeling

For confocal microscopy and FRAP assays, Grb2 variants were covalently labeled using N-hydroxysuccinimide (NHS) ester chemistry. Grb2 Y160E was labeled with [RED-NHS] for initial phase characterization, while for co-recruitment assays, Y160E and WT were labeled with Alexa Fluor 488 and Alexa Fluor 555 (Thermo Fisher Scientific), respectively. Prior to labeling, the protein buffer was exchanged to Labeling Buffer (50 mM HEPES pH 7.4, 300 mM NaCl) to remove primary amines that interfere with the reaction. The coupling reaction was initiated by incubating the protein with a 2-fold molar excess of the reactive dye for 1 hour at room temperature in the dark, following standard bioconjugation protocols [18].

Unreacted fluorophores were removed by Size Exclusion Chromatography (SEC) using a Superdex 75 10/300 GL column. The efficiency of the reaction was quantified by UV-Vis spectroscopy. The Degree of Labeling (DOL) was calculated spectrophotometrically, correcting for the dye’s absorbance at 280 nm according to the correction factors and formulas described by Hermanson [22] and the dye manufacturer’s instructions. Typical DOL values ranged between 0.8 and 1.2. For all imaging experiments, labeled proteins were doped into the unlabeled protein stock at a molar ratio of 1:50 to prevent fluorescence saturation and minimize potential artifacts derived from the fluorophore [18].

### Turbidity Assays

Liquid-liquid phase separation (LLPS) was monitored by measuring solution turbidity (light scattering) at 375 nm using a Multiskan SkyHigh Microplate Spectrophotometer. Experiments were performed in 96-well non-binding surface (NBS) plates (Corning) to minimize surface wetting effects and protein adhesion. Samples were prepared in ice by mixing protein stocks with Assay Buffer and Polyethylene Glycol (PEG) 6000 to reach the indicated final concentrations. The total reaction volume was 100 µL per well. Data was acquired at room temperature (25°C), following standard guidelines for phase separation assays [18].

Phase Diagrams: To map the phase boundaries, turbidity was measured for Grb2 Y160E at concentrations ranging from 20 to 100 µM in the presence of 0–22% (w/v) PEG^6000^. Measurements were performed in triplicate, and data were plotted as the mean absorbance ± standard deviation (SD). Background absorbance from buffer/PEG solutions was subtracted from all readings.

Kinetic Assays: The time-dependent evolution of the condensed phase was monitored for 60–120 minutes. Both WT and Y160E variants were analyzed under fixed crowding conditions (100 µM protein, 20% PEG^6000^). To capture the initial nucleation kinetics, the "dead time" between mixing and the first measurement was minimized to < 30 seconds. Absorbance readings were taken every 90s with 30s shaking before each read to ensure sample homogeneity. Data were normalized to the maximum absorbance value to facilitate kinetic comparison [23].

### Fluorescence and Confocal Microscopy Images

Biomolecular condensates formed by GRB2 WT and Y160E mutation were visualised using wide-field (fluorescence) and confocal microscopies. Protein samples were prepared in 50 mM HEPES buffer (pH adjusted as needed), supplemented with 2–20% (w/v) PEG 6000 (ThermoFisher), and incubated at room temperature for 20 minutes before imaging. A 5 μL droplet of the solution was placed on a 20 × 20 mm coverslip (Deckgläser) for microscopy. Data acquisition was performed using SymPhoTime 64 (PicoQuant), and image analysis was conducted in ImageJ. Confocal fluorescence microscopy was performed using a Leica Stellaris 8 system equipped with a scanning laser and a HyD (Hybrid Detector) module. Samples of Grb2 Y160E and Grb2 WT labelled with Alexa Fluor 488 (green) and Alexa Fluor 555 (orange), respectively, were prepared in 50 mM HEPES buffer at pH 7.4 containing 14% PEG 6000, with each protein at a final concentration of 1 mg/mL. To reduce surface interactions, samples were mounted on mPEG5k-treated silanized 20 × 20 mm coverslips (Deckgläser). Image acquisition was performed using a 63× water-immersion objective (numerical aperture, NA 1.2). Excitation wavelengths were set to 488 nm and 561 nm, with emission collected in the 500–550 nm and 570–620 nm spectral windows, respectively. Three-dimensional reconstructions were obtained by Z-stack scanning at 0.3 μm intervals using a 512 × 512-pixel resolution. Spectral compensation and background subtraction were performed automatically with Leica LAS X software.

### Hyperspectral imaging

Hyperspectral images were acquired using a confocal Leica Stellaris 8 microscope equipped with a HC PL APO CS2 63x/1.40 oil immersion objective (Leica Microsystems). A laser wavelength at 405 nm was used for ACDAN excitation. The ACDAN dissolved in DMSO was directly added to the unlabeled protein solution (100 µM protein concentration in 50 mM HEPES buffer, 14% PEG 6K, pH 7.4) before the experiment, to a final ACDAN concentration of 5 µM. Image acquisition was performed with a frame size of 512 × 512 pixels and a pixel size of 150 nm. The xyλ configuration of the microscope was used, sequentially measuring in 32 channels in the range from 415 to 725 nm. The images were processed using the SimFCS software, developed at the Laboratory of Fluorescence Dynamics, which is available on the webpage (https://www.lfd.uci.edu/globals/ accessed in 2025). The spectral phasor analysis utilises a Fourier transform of the fluorescence emission spectra, from which we calculate two parameters, G and S (as shown in the equations below), that represent the real and imaginary components of the transform.

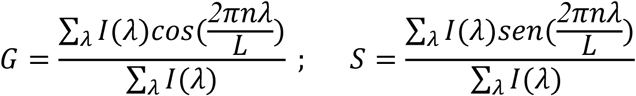

In the equations above, *I(λ)* is the intensity in each step of spectral measurement, *L* is the spectrum range (λ_max_ - λ_min_), and *n* is the harmonic number (=1 in this case). The spectral phasor plot consists of the two parameters G and S plotted in a four-quadrant polar coordinate system. In this case, the wavelength range is captured at each pixel of the hyperspectral image.

### Dynamic Light Scattering (DLS)

DLS measurements were performed using a Malvern Zetasizer Nano ZS instrument equipped with a 633 nm He-Ne laser and operating at a backscattering angle of 173° (NIBS technology). Samples containing 100 µM Grb2 Y160E in Assay Buffer (without PEG) were centrifuged (13.000 rpm, 10 min) and filtered through 0.22 µm PVDF membranes to remove dust and large aggregates. Measurements were conducted in low-volume quartz cuvettes. The hydrodynamic diameter (Dh) was monitored during a controlled temperature ramp cycle (20°C to 5°C to 20°C) with 5°C steps and 120 seconds of equilibration at each point. Data were analyzed using the Zetasizer Software (v7.12) applying the "General Purpose" (non-negative least squares) algorithm to derive the size distribution by intensity, as detailed in Stetefeld et al. [26].

### Fluorescence Recovery After Photobleaching (FRAP)

FRAP experiments were conducted on the Zeiss LSM 880 confocal microscope using the 488 nm argon laser line. A circular Region of Interest (ROI) of 1/2 diameter was defined within the center of a condensate. Following baseline acquisition, the ROI was photobleached using a high-intensity laser pulse (100% power) to reduce fluorescence intensity to <20%. Fluorescence recovery was monitored at low laser power 2% for 120 seconds. Data were normalized to the pre-bleach intensity and corrected for background signal and acquisition photobleaching using a reference non-bleached droplet, following the standardization guidelines by Alberti et al. [18].

### Coarse-Grained Molecular Dynamics Simulations

Simulations were performed using the residue-level COCOMO2 force field [19], which extends the applicability of the original model to accurately capture the phase behavior of systems containing both intrinsically disordered regions and folded domains. In this coarse-grained representation, each amino acid is mapped to a single bead centered at the Cα position. To maintain the structural integrity of the folded SH2 and SH3 domains within the Grb2 monomer (modeled from AlphaFold predictions [21]), secondary and tertiary structures were stabilized using an Elastic Network Model (ENM) applied to residue pairs within the native contact map, as defined by Jussupow et al. [19].

Non-bonded interactions were modeled using a structure-based Lennard-Jones potential for short-range contacts and a Debye-Hückel potential to explicitly account for pH-dependent electrostatic interactions in an implicit solvent environment. Crucially, to correct for the overestimation of interactions in globular domains common in standard CG models, COCOMO2 incorporates a surface exposure scaling factor. This term modulates the interaction strength of each residue based on its solvent-accessible surface area relative to a reference state, ensuring that buried residues in the folded domains contribute less to intermolecular contacts than surface-exposed "stickers" [19].

Simulations were carried out using the OpenMM 8.0.0 toolkit [20]. The system consisted of 250 copies of Grb2 Y160E in a cubic box of 50 nm, corresponding to an effective concentration of approximately 100 µM. The equations of motion were integrated using Langevin dynamics at 298 K with a friction coefficient of 0.01 ps^-1^ and a time step of 20 fs. Production runs were extended for 500 ns to ensure convergence of the self-assembly process. Trajectories were analyzed using MDTraj [27] to calculate cluster size distributions and intermolecular contact maps.

## Supporting information

Support Information

## AUTHOR CONTRIBUTIONS

R.V.R.D. - Conceptualization, Computational Data Investigation, Writing - Original Draft, Data Curation

C. G. - Microscopy experimental Data, writing.

A. S. O. - Expression and Purification of proteins.

G. D. F. - Computational Data Investigation

S. R. T. - Resources, Funding acquisition, Writing - Review & Editing.

A. J. C. F. - Resources, Funding acquisition, Writing - Review & Editing.

F. A. M. - Resources, Funding acquisition, Writing - Review & Editing.

## ACKNOWLEDGMENTS

This work was supported by São Paulo Research Foundation (FAPESP: 2019/24974-0; 2023/01744-5), National Council for Scientific and Technological Development (CNPq: 404982/2025-5). This work received partial financial support from the National Institute of Science and Technology in Innovative Research in Health Sciences – from Nanotechnology to Artificial Intelligence (INCT PICS) sponsored by Brazil’s National Council for Scientific and Technological Development (CNPq), grant no. 408417/2024-2, Coordination of Superior Level Staff Improvement (CAPES), grant no. 88887.197686/2025-00, and São Paulo Research Foundation (FAPESP), grant no. 2025/26818-7. AJCF also thanks FAPESP for support via grants 2024/17967-6, 2024/17969-9, 2023/04532-9, and CNPq via grant 309476/2025-9.

We also thank the National Laboratory for Scientific Computing (LNCC/MCTI, Brazil) for providing HPC resources for the SDumont supercomputer (Project prodex-grb2) URL: http://sdumont.lncc.br.

## DECLARATION OF GENERATIVE AI AND AI-ASSISTED TECHNOLOGIES IN THE WRITING PROCESS

During the preparation of this work the authors used ChatGPT in order to improve writing fluency. After using this tool, the authors reviewed and edited the content as needed and took full responsibility for the content of the publication.

## Support Information

**Supplementary Figure S1.**
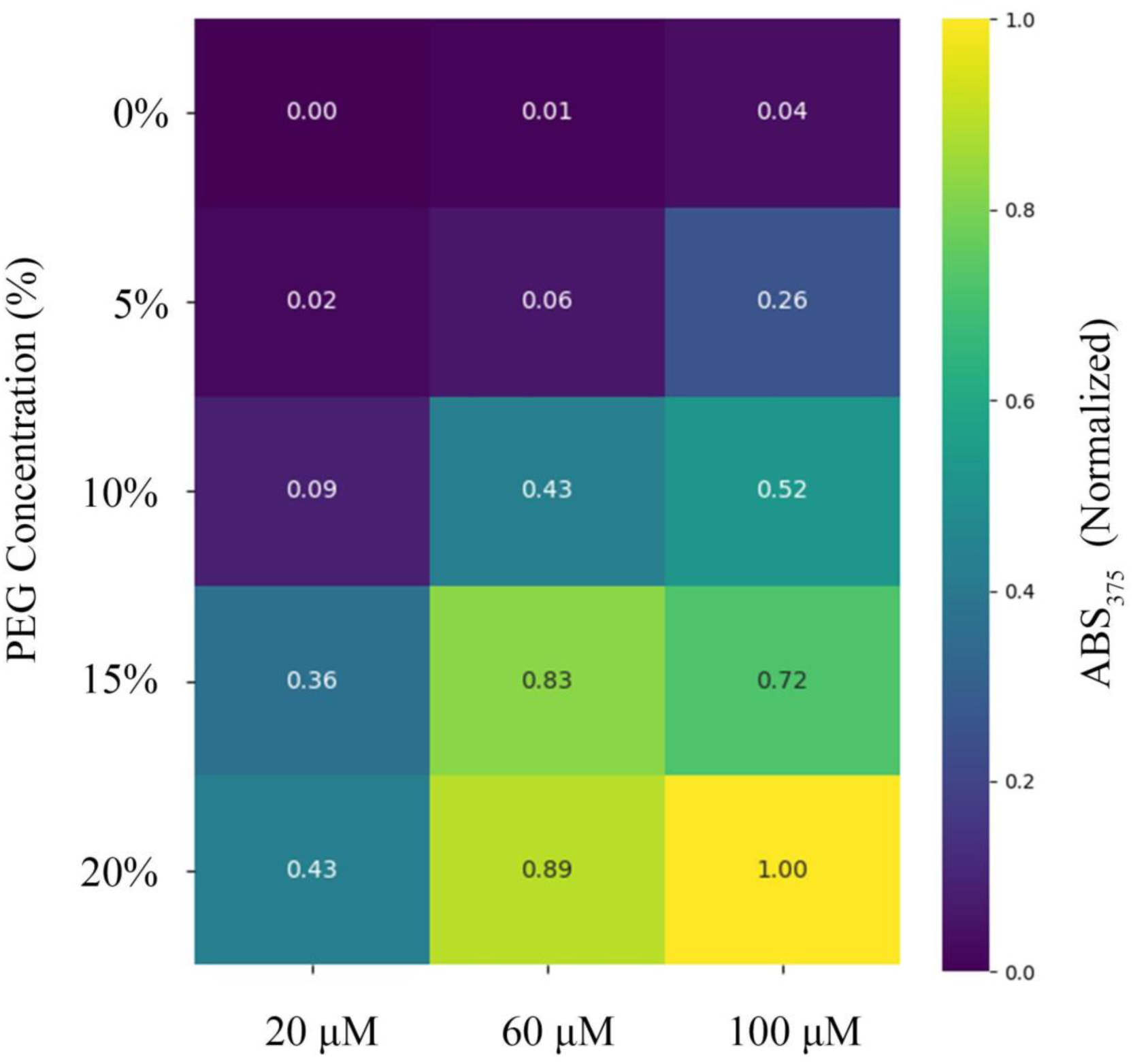
Detailed turbidity heatmap of Grb2 Y160E phase separation. The phase diagram was mapped by measuring solution turbidity (Absorbance at 375 nm) across a grid of protein concentrations (20, 60, and 100 µM) and crowding agent concentrations (0–20% PEG-6000). The heatmap colors represent normalized turbidity values, ranging from **0.0 (dark purple)**, indicating a dispersed monomeric phase, to **1.0 (bright yellow)**, indicating maximal condensation. The numbers within each cell represent the specific normalized absorbance value for that condition. The data illustrate a clear phase transition boundary where condensation is triggered only above a critical threshold of protein and crowder concentrations.

**Figure S2.**
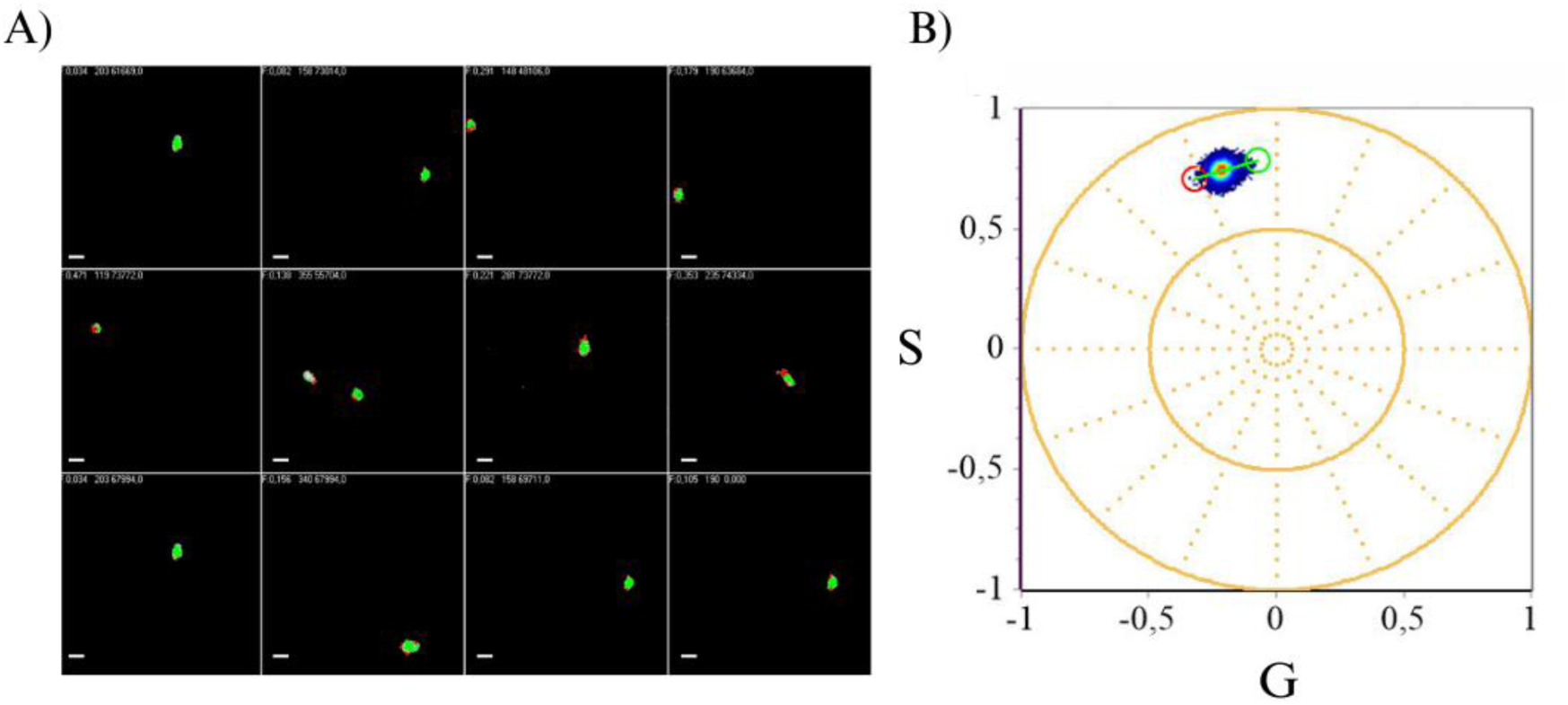
Hyperspectral imaging of ACDAN reports the local microenvironment within Grb2 condensates. (A) Representative phasor coordinate maps generated from ACDAN hyperspectral datasets, showing the spatial distribution of pixels assigned to the selected regions of the phasor plot. Scale bars 2,5 um. (B) Corresponding spectral phasor plot (G–S) displaying a single compact population; the colored cursors denote the regions used to generate the maps in (A).

**Supplementary Video S3. Spontaneous self-assembly and condensate formation of Grb2 Y160E *in silico*.** The video displays the Coarse-Grained Molecular Dynamics (CG-MD) trajectory of 250 Grb2 Y160E monomers over a 500 ns simulation. The system initially starts from a dispersed, homogeneous state. As the simulation progresses, the monomers undergo rapid nucleation to form transient clusters, which subsequently coalesce into a single, dense, and stable mature condensate. To highlight the dynamic, multichain nature of the assembly, each individual protein chain is assigned a distinct color. The proteins are rendered in a "tube" format, reflecting the Ca-only resolution of the coarse-grained model. The trajectory was rendered against a black background for enhanced visual contrast, and Periodic Boundary Conditions (PBC) were wrapped to ensure the continuous visualization of the clusters across the simulation box boundaries. Video rendered using Visual Molecular Dynamics (VMD) software.

