## Supplementary material for "Beyond signaling activation: Phosphorylation modulates Grb2 phase separation to create multivalent scaffolds": Support Information

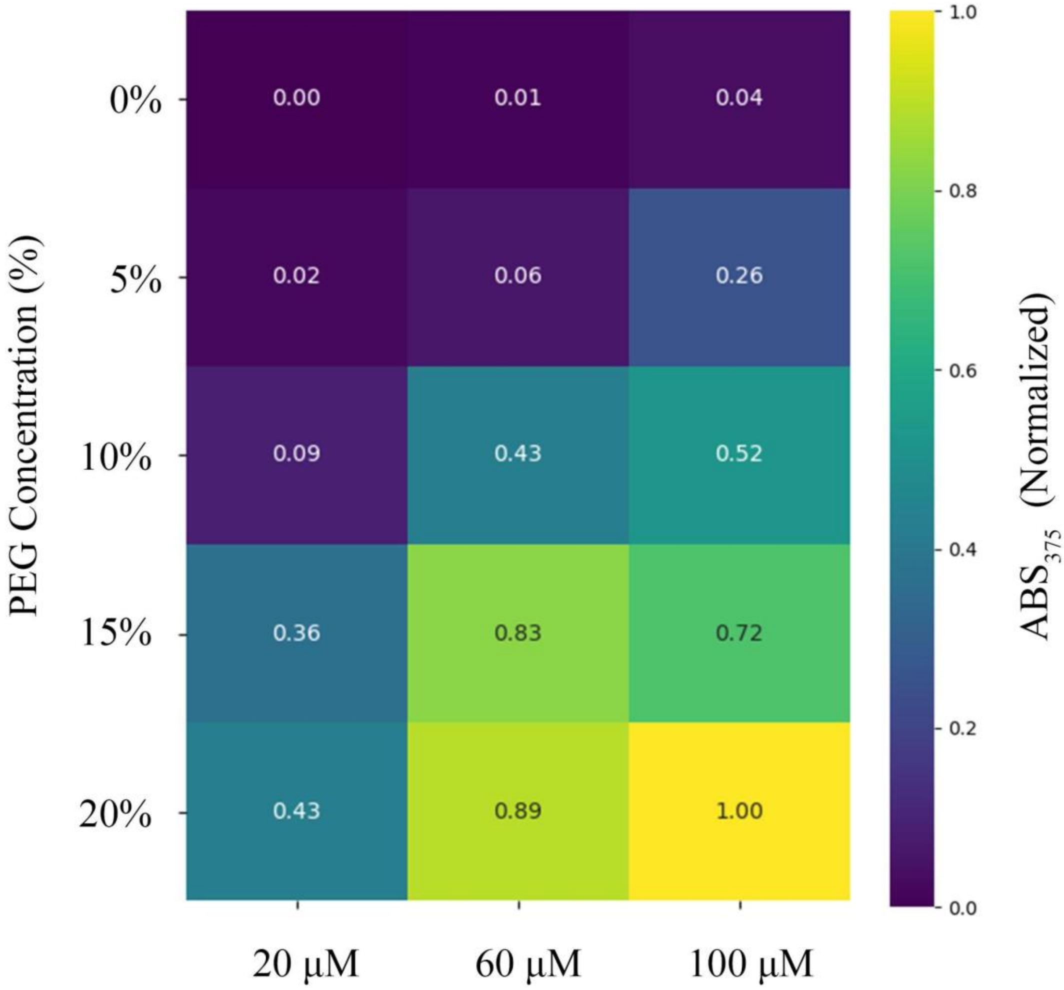


**Supplementary Figure S1. Detailed turbidity heatmap of Grb2 Y160E phase separation.** The phase diagram was mapped by measuring solution turbidity (Absorbance at 375 nm) across a grid of protein concentrations (20, 60, and 100 µM) and crowding agent concentrations (0–20% PEG-6000). The heatmap colors represent normalized turbidity values, ranging from **0.0 (dark purple)**, indicating a dispersed monomeric phase, to **1.0 (bright yellow)**, indicating maximal condensation. The numbers within each cell represent the specific normalized absorbance value for that condition. The data illustrate a clear phase transition boundary where condensation is triggered only above a critical threshold of protein and crowder concentrations.

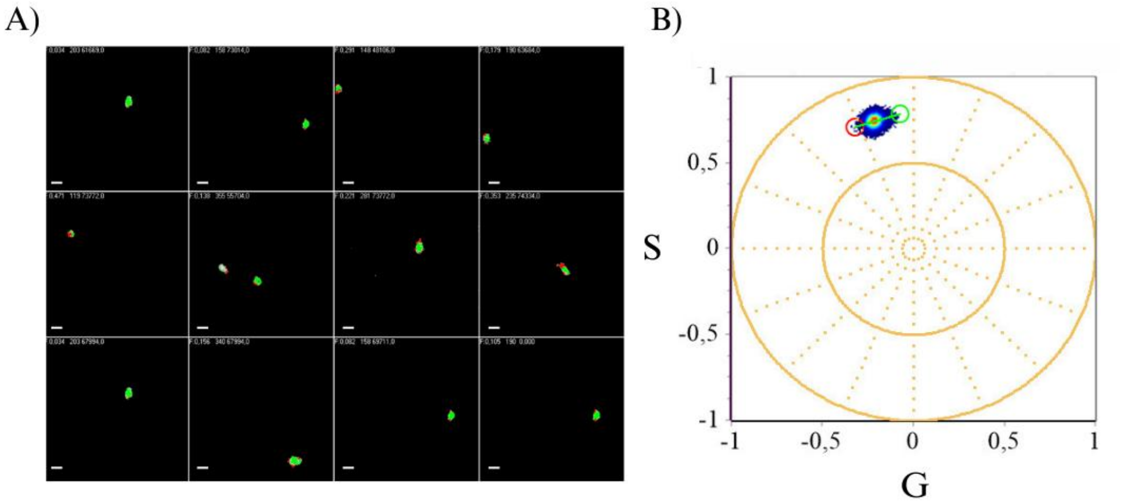


**Figure S2. Hyperspectral imaging of ACDAN reports the local microenvironment within Grb2 condensates.** (A) Representative phasor coordinate maps generated from ACDAN hyperspectral datasets, showing the spatial distribution of pixels assigned to the selected regions of the phasor plot. Scale bars 2,5 um. (B) Corresponding spectral phasor plot (G–S) displaying a single compact population; the colored cursors denote the regions used to generate the maps in (A).

**Supplementary Video S3. Spontaneous self-assembly and condensate formation of Grb2 Y160E *in silico*.** The video displays the Coarse-Grained Molecular Dynamics (CG-MD) trajectory of 250 Grb2 Y160E monomers over a 500 ns simulation. The system initially starts from a dispersed, homogeneous state. As the simulation progresses, the monomers undergo rapid nucleation to form transient clusters, which subsequently coalesce into a single, dense, and stable mature condensate. To highlight the dynamic, multichain nature of the assembly, each individual protein chain is assigned a distinct color. The proteins are rendered in a "tube" format, reflecting the Ca-only resolution of the coarse-grained model. The trajectory was rendered against a black background for enhanced visual contrast, and Periodic Boundary Conditions (PBC) were wrapped to ensure the continuous visualization of the clusters across the simulation box boundaries. Video rendered using Visual Molecular Dynamics (VMD) software.
